# Cell-type-specific Labeling of Endogenous Proteins Using the Split GFP System in *Drosophila*

**DOI:** 10.1101/2024.05.06.592806

**Authors:** Melissa Ana Inal, Kota Banzai, Rie Kamiyama, Daichi Kamiyama

## Abstract

Accurate identification of the locations of endogenous proteins is crucial for understanding their functions in tissues and cells. However, achieving precise cell-type-specific labeling of proteins has been challenging *in vivo*. A notable solution to this challenge is the self-complementing split green fluorescent protein (GFP_1-10/11_) system. In this paper, we present a detailed protocol for labeling endogenous proteins in a cell-type-specific manner using the GFP_1-10/11_ system in fruit flies. This approach depends on the reconstitution of the GFP_1-10_ and GFP_11_ fragments, creating a fluorescence signal. We insert the *GFP*_*11*_ fragment into a specific genomic locus while expressing its counterpart, *GFP*_*1-10*_, through an available Gal4 driver line. The unique aspect of this system is that neither GFP_1-10_ nor GFP_11_ alone emits fluorescence, enabling the precise detection of protein localization only in Gal4-positive cells expressing the GFP_11_ tagged endogenous protein. We illustrate this technique using the adhesion molecule gene *teneurin-m* (*Ten-m*) as a model, highlighting the generation and validation of GFP_11_ protein trap lines via Minos-mediated integration cassette (MiMIC) insertion. Furthermore, we demonstrate the cell-type-specific labeling of Ten-m proteins in the larval brains of fruit flies. This method significantly enhances our ability to image endogenous protein localization patterns in a cell-type-specific manner and is adaptable to various model organisms beyond fruit flies.

## 1. Introduction

Fluorescent proteins (FPs) have become a cornerstone in cell and developmental biology. Observing the behaviors of proteins tagged with FPs in live specimens provides critical insights into various biological processes, including macromolecular assembly, protein trafficking, and protein translation [1]. Typically, these experiments involve overexpressing the protein of interest in cells or tissues of model organisms.

While this overexpression strategy is highly effective, it can lead to artifacts due to the use of basal or tissue-specific promotors, often differing from the transcriptional regulation of target genes. These artifacts include protein mislocalization and aggregation, abnormal organelle morphology, and the mistiming of cellular signaling events [2, 3]. To mitigate these issues, we employ endogenous gene tagging, where an FP’s coding sequence is inserted into a specific genomic locus [4]. This allows the FP-fused proteins to be expressed under native transcriptional and translational controls.

Historically, full-length FPs have been widely used in various applications. More recently, the split GFP_1-10/11_ system has emerged as a valuable tool for live-cell protein imaging, especially with *in situ* imaging. Furthermore, due to its binary nature reconstitution can be achieved in a cell-specific manner [5-9]. In this system, the super-folder GFP sequence is divided between β-strand 10 and 11 (GFP_1-10_ and GFP_11_)[10].

Neither GFP_1-10_ nor GFP_11_ alone is fluorescent. The GFP_11_ fragment, a 16-residue peptide, only when in the presence of its counterpart, GFP_1-10_ can self-assemble with each other to reconstitute a functional GFP. Our previous research has used this system to investigate the adhesion molecule Ten-m in Drosophila, previously known for its expression in the nervous system but whose synaptic functions required further exploration [11, 12]. By using a specific MiMIC (Minos Mediated integration cassette) insertion line that contains a *MiMIC* fragment, we have replaced the Swappable Integration Cassette (SIC) provided in the Minos backbone with the *GFP*_*11*_ fragment through Recombinase-Mediated Cassette Exchange (RMCE). This replacement has enabled the creation of *ten-m* protein-trap lines tagged with *GFP*_*11*_, allowing us to visualize the endogenous localization of Ten-m in the central nervous system (CNS). An additional advantage of this approach is the tandem repeating capability of GFP_11_ due to its small size, enabling the tagging of a gene with multiple GFP_11_ repeats (e.g., GFP_11×7_) for enhanced fluorescent signals. Using this technique, we enhanced the Ten-m signal and achieved cell-type-specific labeling in various neuron types by expressing GFP_1-10_ in specific cells of the Drosophila brain.

In this chapter, we present a detailed protocol for this innovative protein-tagging approach, focusing on its application to the *ten-m* gene locus in *Drosophila*. We detail the procedures for RMCE line generation via MiMIC insertions, including the creation and injection of the plasmid and the subsequent crossing schemes. We emphasize the importance of confirming the directionality of GFP_11_ insertion for successful protein labeling. Lastly, we discuss the process for achieving cell-type-specific labeling of Ten-m proteins in the larval brains of fruit flies. This method can be adapted to label a wide range of proteins in a cell-type-specific manner.

## 2. Materials

1. **DNA Preparation**
  - Modified GFP11 plasmid in phase 1(RRID: Addgene_172068)
  - Modified GFP11×7 plasmid in phase 1 (RRID: Addgene_172067)
  - Nano spectrophotometer for measuring DNA concentration (Implen)
2. **Injection**
  - MiMIC gene-trap flies of interest: *y[1] w[*]; Mi{y[+mDint2] =MIC}Ten-m[MI07828]* (RRID: BDSC_44887)
  - nanos-ΦC31 integrase stock (RRID: BDSC_ 34771)
  - Fly cage (Genesee Scientific #59-100)
  - Standard grape juice plates
  - *GFP*_*11*_ and *GFP*_*11×7*_ plasmids from DNA preparation step in the appropriate reading frame (RRID: Addgene_172068, Addgene_172067). Ten-m is found in phase 1 thus it is possible to use the AddGene plasmids for *GFP*_*11*_ insertions.
  - Capillary glass with filament (Harvard Apparatus #30-0032)
  - Needle puller (Narishige #PN-31)
  - Spin filtration column (Co-Star #8160)
  - 18 mm x 18mm cover glass (Corning #2845-18)
  - Glass slide (Eisco #BI0082)
  - Needle holder (Narishige H-7)
  - Upright microscope with LED light source (Nikon Eclipse Ci)
  - 100% Ethanol (Sigma #E7023)
  - Mineral oil (source)
  - FemtoJet 5247 Microinjector (Eppendorf, Discontinued)
  - Dumont #5 forceps (Fine Science Tools #11252–20)
  - Standard food vial
3. **Line Production and Validation**
  - Balancer adult flies (*y[1]w[*]; TM3, Sb[1]/TM6B, Tb[+]*, RRID: BDSC_3720)
  - Microcentrifuge tubes
  - PureLink™ Genomic DNA Mini Kit (Invitrogen, K1820-01)
  - GFP_11_ Tag-specific primers: GFP11.for (CGTGACCACATGGTCCTTCATGAGT) and GFP_11_.rev (TGTAATCCCAGCAGCATTTA). See Note 3 for GFP_11×7_ primer sequences to use in validation.
  - MiMIC-specific primers: MiL-F (GCGTAAGCTACCTTAATCTCAAGAAGAG). See Note 4 for MiL-R sequence needed for detailed orientation validation.
  - For PCR reaction: Taq 2x Master Mix (New England Biolabs Inc, M0270L).
4. **Tissue-specific Reconstitution**
  - GFP_11_ RMCE line (generated in the previous step)
  - Gal4 driver for cell-type of interest: *elav-Gal4* (RRID: BDSC_8765 or BDSC_8760), *dlip2-Gal4*
  - (RRID: BDSC_35716), *eve*-Gal4 (RRID: BDSC_7473), *ato-Gal4* (RRID: BDSC_9494)
  - *UAS-GFP*_*1-10*_^*sec*^ (for extracellular GFP_11_; RRID: BDSC_93190) or *UAS-GFP*_*1-10*_ (for intracellular GFP_11_; RRID: BDSC_93189)
5. **Imaging**
  - 100 × 1.45 NA oil immersion objective (Nikon).
  - Inverted fluorescence microscope (Nikon) with a Dragonfly Spinning disk confocal unit (Andor). Other confocal microscopes equipped with a 488-nm laser are suitable.
  - Glass slide
  - 22×22 mm square cover glass, #1.5 (Corning)
  - Mounting media (1x PBS)
  - Fiji software (NIH). Any other image processing software can be used.

## 3. Methods

### 3.1 Pre-experiment Reagent and Fly Preparation

#### 3.1.1 Preparation of DNA for Transgenic Flies

The MiMIC insertion occurs randomly within the genome. The MiMIC insertion site, in relation to the open reading frame of the target gene, is a crucial consideration in selecting the appropriate SIC vector for RMCE, ensuring the correct translation of the gene product without a frameshift. There are 3 phases of RMCE plasmids (Phase 0, 1, and 2) which needs to be matched to the relative position of the open reading frame of a MiMIC insertion. Our gene of interest in this chapter, *ten-m* found in Phase 1, has been successfully tagged with *GFP*_*11*_ by insertion into the RMCE Plasmid Phase 1 (RRID: Addgene_172068). If *GFP*_*11×7*_ will be used for tagging other genes in phase 1, the modified plasmid is available through Addgene (RRID: Addgene_172067). Otherwise, it is possible to generate the *GFP*_*11×7*_ plasmid by standard cloning techniques their respective RMCE plasmids as previously published. For this step we sourced GFP_11_ from an oligo synthesizer company (i.e., Integrated DNA Technologies) and cloned into the RMCE plasmids, ensuring 100 ng/μl final concentration of DNA for the subsequent injection step. See Figure 1 for plasmid design.

**Figure 1.**
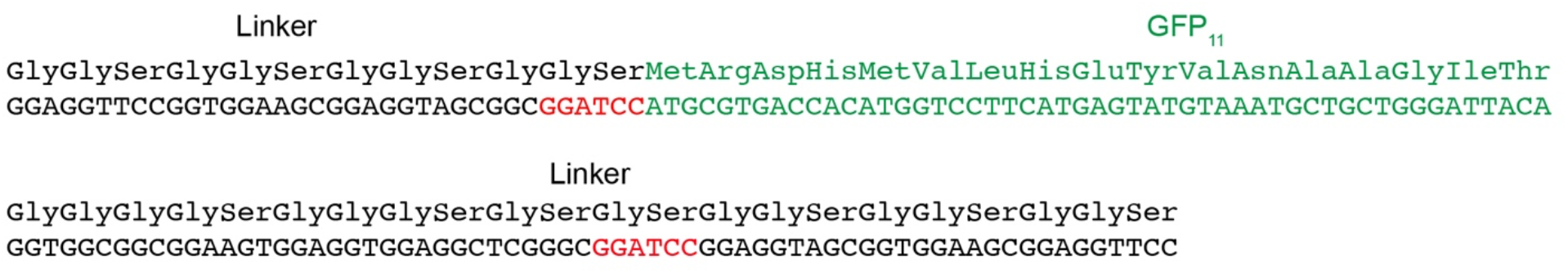
Diagram of the GFP_11_ with Linker Sequence for RMCE Plasmid. This illustrates the specific insertion of the *GFP*_*11*_ coding sequence (green) into an established protein-trap plasmid. The insertion was facilitated using the *BamHI* sites (red) located between the peptide linkers. These peptide linkers, composed of a series of Glycine-Glycine-Serine amino acid residues, provide flexibility between the GFP_11_ fragment and the protein being tagged.

#### 3.1.2 Fly Embryo Preparation and Needle Preparation for DNA Injection

For proper integration of external DNA (such as the *GFP*_*11*_ fragment) into the fly genome, it is essential to prepare the appropriate progeny before injection. For RMCE, the flies must carry the phiC31 integrase in their germline via nanos to allow for the *GFP*_*11*_ cassette to exchange the MiMIC SIC at the phiC31 attP sites. This step includes setting up a genetic cross and training the adult flies to lay eggs.

1. Start preparation 2-3 days prior to the experiment by setting up a fly cage with grape juice plates. Cross the virgin females carrying the phiC31 integrase on the X chromosome to the males carrying the MiMIC insertion in the Ten-m locus on the third chromosome. Each cage should maintain a male-to-female ratio of 1:5, encompassing a total of 100 adult flies per cage.
2. Transfer the flies into a cage and cover it with a grape juice plate. Maintain by changing the plate each day with a new grape juice plate with a smear of fresh yeast paste at 25°C (or room temperature) for approximately 3 days to allow the flies to synchronously lay eggs.

### 3.2 RMCE Experiment

#### 3.2.1 Injection of Plasmid into Flies

Although this step can be outsourced to various companies (e.g., BestGene, Inc. or Rainbow Transgenic Flies, Inc.), here we describe the methods to generate a transgenic line in a lab setting that has an injection arrangement. The primary consideration is the proper execution of the injection process to maximize the number of survivors after injection. This is crucial for yielding ‘founders’ that transmit the GFP_11_ cassette to the next generation via germline transmission. Typically, more than 1-10% of the embryos become adults after injection. Our protocol below is based on classic protocols with minor modifications [13].

1. Pull needles using a Narishige needle puller set to a heater level of 81.6 and a magnet level of 45.5, based on our puller settings. Note that this setup may vary depending on your specific puller. The guidelines provided by the Kaufman lab offer recommendations for needle shapes suitable for injecting fly eggs [14].
2. Get the DNA ready for injection by passing it through a spin filtration column to prevent the needle from clogging up.
3. Load 1.5 μl of DNA per needle using a micro-loader pipette tip.
4. 2 hours prior to injection, replace the grape juice plate in the cage and collect embryos every hour. This allows for synchronization of embryo development, ensuring that over 90% of the embryos are at stage 2, which is before the syncytium stage.
5. Flush the embryos into a cell strainer (∼40 μm strainers would suit this purpose) with deionized (DI) water.
6. Transfer them gently using a paintbrush from the cell strainer and place them on an 18mm x 18mm cover glass.
7. With the paintbrush, align approximately 80 embryos horizontally on a cover glass, ensuring they are oriented with the same anterior-posterior axis, as the needle is inserted only from the posterior. Caution: Maintain a moist condition by keeping the brush tip and embryos wet with DI water.
8. Attach the cover glass onto a glass slide and then place onto the microscope stage.
9. Allow the embryos to dry for 1-2 minutes to ensure they firmly attach to the cover glass.
10. Attach the pulled needle from step 1 to a needle holder and position the needle tip near the posterior end of the embryos under the microscope.
11. Cover the line of embryos with 100% ethanol. Note: We use ethanol instead of halocarbon oil for a better survival rate under our experimental conditions (this may vary from lab to lab).
12. Set the injection pressure (Pi) to 300 hectopascals, the injection time (Ti) to 0.1 seconds, and the compensation pressure (Pc) to 300 hectopascals on the Microinjector FemtoJet.
13. Insert the needle tip into the posterior fifth of each embryo, where the germline is located, and inject the loaded plasmid into the embryos.
14. Carefully take out the cover glass from the microscope and transfer it to a food vial.
15. Repeat steps 7 to 14 until you have injected up to 1,000 embryos. Place each cover glass in a separate vial.
16. Keep the vials at 25°C. The embryos will take approximately 9-10 days to develop into adults.

#### 3.2.2 Establishing Stable Transgenic Fly Stock

Once DNA injection leads to adult flies, identifying the RMCE progeny is crucial for establishing a stable transgenic line, a process aided by tracking the presence or absence of yellow^+^ which is a constituent of MiMIC that will be excised upon successful recombination. Typically, approximately 10% of the injected adults become founders, producing progeny with a yellow-body color. This progeny can then be established as a stable line for studying GFP reconstitution in various tissue types.

1. Collect the resulting adults from section 3.2.1 and cross individual flies with an appropriate balancer for the third chromosome such as *yw; TM3, Sb[1]/TM6B, Tb[+]*. (See Note 2 for establishing stable lines)
2. Collect single transgenic male flies with the absence of the yellow marker that has a balancer or a dominantly marked chromosome and cross with the balancer virgin females.
3. Sibling-cross the balanced flies to establish stock. See Figure 3 for an example crossing scheme.

### 3.3 Post-genome Editing Analysis

#### 3.3.1 PCR Analysis for Validation of RMCE Event

After selecting progeny with a yellow-body color, we validate the successful RMCE event and the correct orientation of the *GFP*_*11*_ insertion through PCR analysis. By using standard DNA extraction and PCR reactions, we can assess the presence and orientation of the insert prior to *in vivo* testing. It is important to note that after RMCE, the *GFP*_*11*_ fragment can be inserted in either direction – 5’ to 3’ or 3’ to 5’ -thus, validation of the orientation is required.

1. Using the manufacturer’s protocol, extract DNA of 5 adult transgenic flies from F1 generation (PureLink™ Genomic DNA Mini Kit, Invitrogen). While we used the same volumes and concentrations of the reagents, we find that elution with 20 μl of Elution Buffer is sufficient in the final step.
2. Next, for the PCR reaction prepare PCR master mix with 10 μl reaction volume for each candidate. See Materials for specific primer sequences used.
  a. Master mix recipe: 1 μl DNA, 5 μl Taq 2x Master Mix, 3 μl milliQ water, 0.5 μl per primer in each reaction listed.
    i. MiL-F and GFP_11_.for
    ii. MiL-F and GFP_11_.rev
3. Set the cycling conditions as follows:
  a. Initial denaturation at 95° for 30 seconds
  b. 35 cycles at 94° for 30 seconds, 67° for 1 minute, and 68° for 30 seconds
  c. Final extension at 68° for 5 minutes
  d. Hold at 10 ºC.
4. Run PCR products on agarose gel. See Figure 2 for an example PCR validation for assessment of orientation (See Note 4 for detailed interpretation of results)

**Figure 2.**
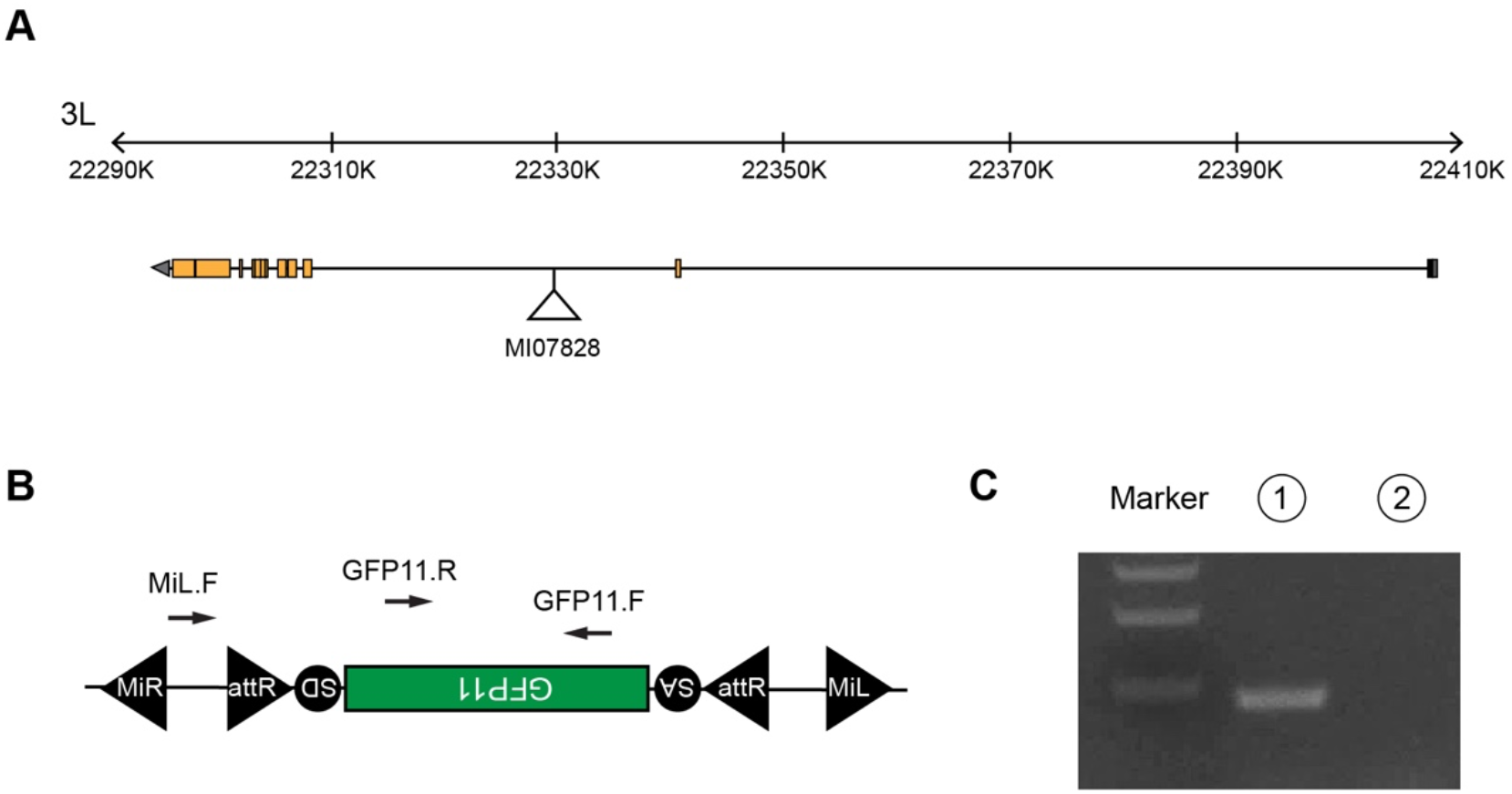
Assessment of GFP_11_ Insertion Orientation in the Ten-m MiMIC Insertion. (**A**) Schematic of a MiMIC insertion (MI07828) in the *ten-m* gene (orange boxes denote exon regions) (**B**) Representation of the RMCE process with *attP* sites, leading to the substitution of the target sequence with the *GFP*_*11*_ sequence from our plasmid construct. This process allows for the integration of *GFP*_*11*_ in two possible orientations, with the figure showing the preferred orientation. SA (splice acceptor) and SD (splice donor). This diagram also highlights the use of three specific primers for PCR reactions, which enable the determination of the GFP_11_ integration orientation. (**C**) An agarose gel image displays the PCR products, indicating that *GFP*_*11*_ has been inserted in the desired orientation. The numbers above the gel lanes correspond to the primer pairs used: 1 - MiL.F and GFP_11_.for; 2 - MiL.F and GFP_11_.rev.

**Figure 3.**
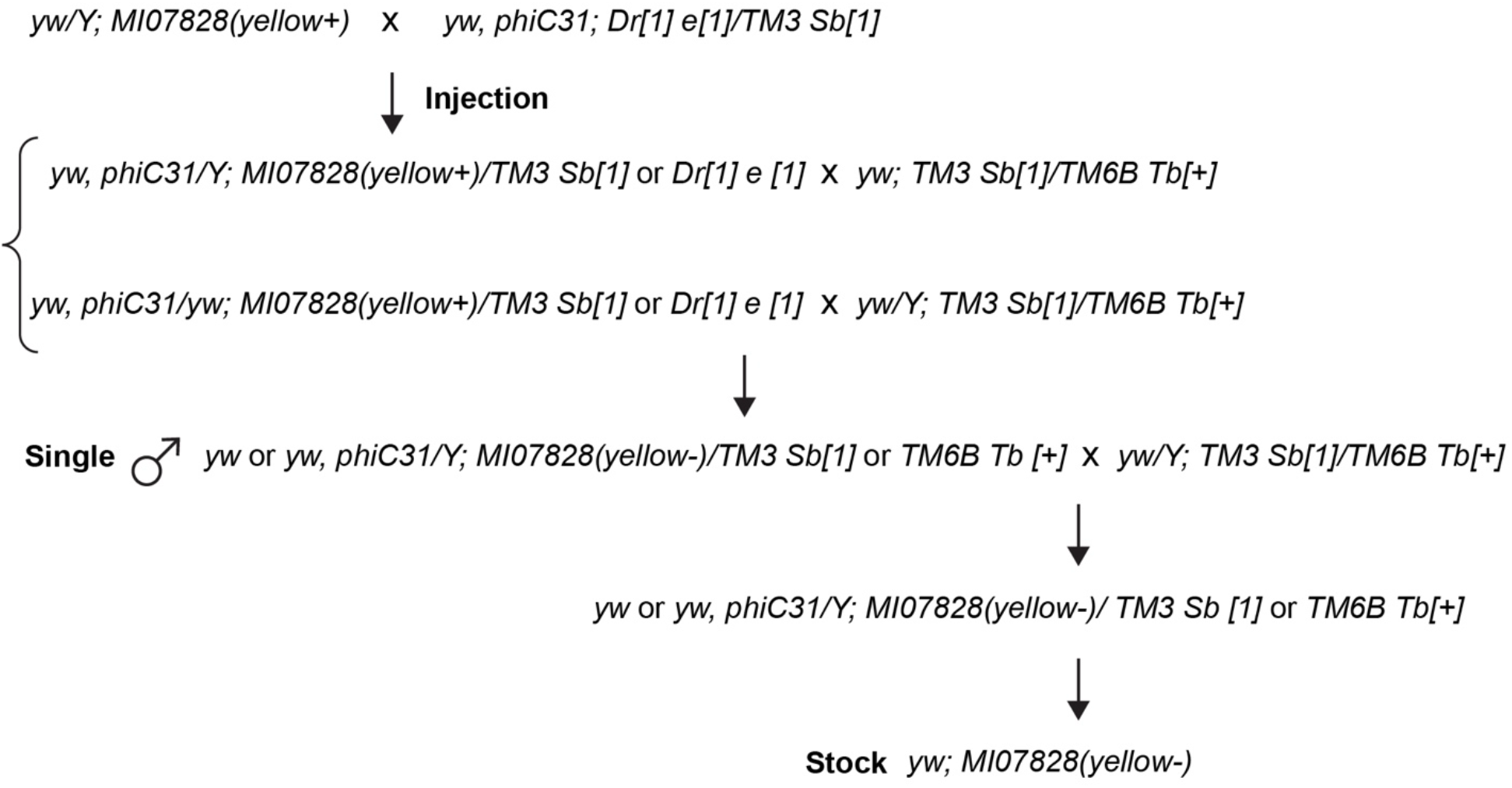
Crossing Schemes for GFP_11_ Line via MiMIC Insertion Using RMCE. This figure illustrates the specific crosses performed to establish the *GFP*_*11*_ line.

#### 3.2.2 Protein Labeling in Specific Cell Types within Flies

The functional integration of *GFP*_*11*_ into a specific genomic locus and the reconstitution of those split GFP fragments can be validated using fluorescence microscopy. Further, increasing the number of *GFP*_*11*_ repeats can be tested for localizing low expression level proteins.

1. To achieve GFP reconstitution in a cell-specific manner, the UAS-Gal4 system can be used, where the GFP_1-10_ component will be expressed under tissue specific Gal4 activity such as *elav*-Gal4. If the localization pattern of the targeted protein is unknown, a basal promoter Gal4 driver such as *Ubi*-Gal4 (RRID: BDSC_32551) can be used. Depending on the GFP11×7 insertion site on the target protein, cytosolic (RRID: BDSC_93189) or secreted GFP_1-10_ (RRID: BDSC_93190) should be used. This step can be done in parallel to the transgenic fly generation (section 3.3), to save time.
2. Using this stable line, cross it to the GFP_11_ RMCE line generated to visualize reconstituted GFP_1-10/11_ signal to track protein localization.
3. Age the progeny of these flies to desired developmental stage (e.g., 2^nd^ or 3^rd^ instar larvae).
4. Dissect larval brains or the tissue of interest, if necessary.
5. Mount the tissue onto a clean glass slide using appropriate mounting media (e.g., 1x PBS).
6. Using a confocal microscope with an appropriate objective lens (e.g. 100x), acquire the image of reconstituted GFP_1-10/11_ with a 488 nm laser.
7. If the reconstitution signal is weak due to a single copy of the *GFP*_*11*_, it is possible to amplify the signal by generating a transgenic fly with the *GFP*_*11×7*_ construct by repeating sections 3.1-3.5. See Figure 4 for comparison of *GFP*_*11*_ and *GFP*_*11×7*_ tags in the larval brain.
8. Use ImageJ or other image analysis software to analyze the localization data. See Figure 5 for a representative confocal image endogenous Ten-m labeled with split GFP at 100x magnification in different cell-types.

**Figure 4.**
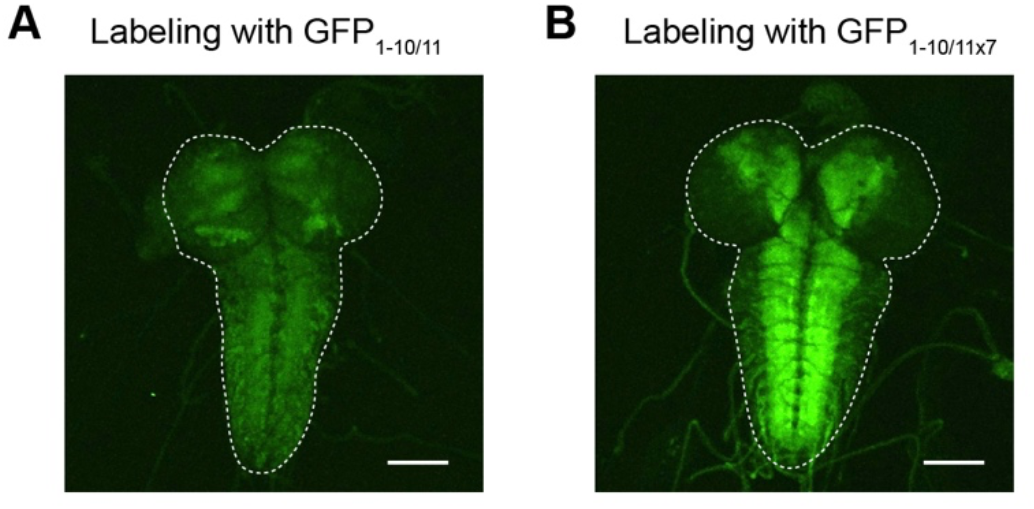
Reconstituted GFP Signals from GFP_11×1_ and GFP_11×7_. Representative images display the reconstituted signals from GFP_11_ (A) and GFP_11×7_ (B). To visualize Ten-m, we crossed either the *GFP*_*11*_ or the *GFP*_*11 × 7*_ strain with a pan neuronal *GFP*_*1–10*_ expression line. Scale bars: 100 μm.

**Figure 5.**
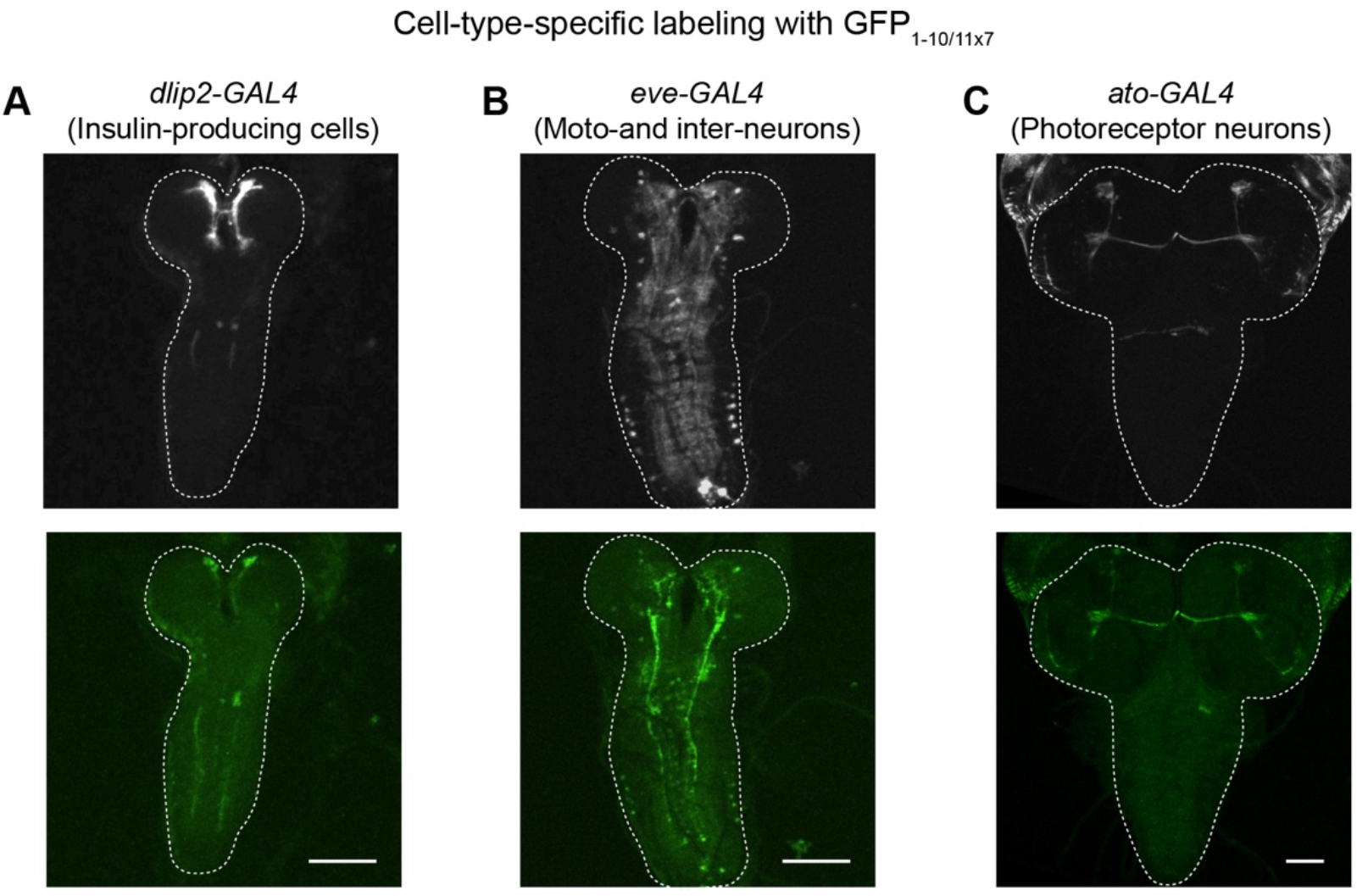
Cell-Type–Specific Labeling of Ten-m Proteins. Representative images demonstrate cell-type– specific fluorescence of a CD4 membrane marker (Top) and Ten-m (Bottom). The *GFP*_*11 × 7*_ protein trap line of ten-m was crossed with various *GFP*_*1–10*_ expression lines. The following Gal4 lines were used to drive *UAS-GFP*_*1–10*_ expression: (A) *dlip2-Gal4*, (B) *eve-Gal4*, and (C) *ato-Gal4*. Dotted lines mark the boundaries of the larval brains. Scale bars: 100 μm.

## 4. Notes

1. The protocol we describe takes advantage of the MiMIC insertion collection generated as a part of the Gene Disruption Project (GDP) which allows for RMCE [15-17]. There are 17,500 MiMIC collection lines reported to be generated as part of the GDP. Of these lines, 1,860 of them have the MiMIC insertion in the intron positioned between the coding exons (also known as coding intron). These are the insertions that can be used to trap proteins with genetic tags such as the *GFP*_*11*_ [7] or a full-length *GFP* cassette as previously reported [15].

Since *Minos* integrates randomly into the genome, it is important to consider where MiMIC is inserted as it may disrupt the gene function and prevent normal protein localization. According to the pilot studies, the GFP-tagged proteins through RMCE into a coding intron does not disrupt gene function in ∼75% of the cases, thus allowing studies of endogenous localization [16, 17]. To assess whether protein function or localization is affected in these lines, a complementation experiment or immunostaining experiment may be needed. The MiMIC fly stock information is maintained under the GDP website. To select a target protein, the information about the type of insertion site whether in the coding intron or elsewhere, as well as information for ordering flies through the BDSC can be found through the following link: https://flypush.research.bcm.edu/pscreen/mimic.html. Additionally, CRIMIC lines, also generated by the Bellen Lab, provide an additional library to utilize for RMCE based GFP_11_ tagging of endogenous genes [18-20]. If the genes of interest are not available in either library for protein trapping, CRISPR-mediated knock-in strategies can be incorporated to precisely modify target proteins, expanding the range of experimental possibilities [21].

1. It is recommended to screen at least 10 independent candidates for 5 positive RMCE events which will guarantee 2-3 lines with proper insertion orientation.
2. For validation of the GFP_11×7_ construct insertion in section 3.2.1, we used the following primers: GFP_11×7_.for (CGTGACCACATGGTCCTTCATGAGTATGTAAATGCTGCTGGGATTACA) and GFP_11×7_.rev (GGTGATACCGGCAGCATTGACATATTCGTGCAGGACCATAT) as the appropriate primer sequences.
3. If gene and original MiMIC insertion are in the same orientation, then the successful RMCE should screen positive for PCR reactions 2 only. If gene and original MiMIC insertion are in opposite orientations, then the successful RMCE should screen positive for PCR reactions 1 only. For more detailed validation PCR reactions should also be run with the MiL-R (CGCGGCGTAATGTGATTTACTATCATAC) primer. More thorough description of such approach can be found in the original manuscript [15].

## 5. Concluding remarks

A collection of approximately 1,800 MiMIC insertions, which are inserted in coding introns of specific genes, is now available [16, 17]. This resource enables the potential creation of a vast library of GFP_11_-tagged genes, facilitating the study of the functions of proteins encoded by these genes. The generation of this library opens a new era in *in vivo* proteomic analysis, overcoming spatiotemporal limitations of existing approaches [22]. In this chapter, we detailed the protocol for creating a RMCE line of *GFP*_*11*_, using the genomic locus of Ten-m as an example. We demonstrated cell-type-specific labeling using various *GFP*_*1-10*_ expression lines. These proteins can be imaged in unfixed tissues and cells. Furthermore, by combining other colors of split FP systems, it is possible to investigate the interactions between Ten-m and other molecules [7]. This protein-labeling strategy offers a new capability to study protein behaviors with unprecedented resolution in vivo and is likely to be extended to other organisms, including vertebrates.

## 6. Acknowledgments

We thank Miyuki Ando Fitch for her technical assistance and the Bloomington Drosophila Stock Center for providing the fly strains. We are also grateful to all members of the Kamiyama laboratory for their critical reading of the manuscript. This work was supported by NIH grants R01 NS107558 and R21 NS128750.

